# Dextranol: A better lyoprotectant

**DOI:** 10.1101/490441

**Authors:** Bryan Jones, Advitiya Mahajan, Alptekin Aksan

**Affiliations:** Biostabilization Laboratory, Department of Mechanical Engineering, University of Minnesota, Minneapolis, Minnesota, United States of America

## Abstract

Dextranol, a reduced dextran, prevents damage to stored dry protein samples that unmodified dextran would otherwise cause. Lyoprotectants like the polysaccharide dextran are critical for preserving dried protein samples by forming rigid a glass that protects entrapped protein molecules. Stably dried proteins are important for maintaining critical information in clinical samples like blood serum. However, we found that dextran reacts with serum proteins during storage, producing high-molecular weight Amadori-product conjugates. These conjugates appeared in a matter of days or weeks when stored at elevated temperatures (37° or 45°C), but also appeared on a timescale of months when stored at room temperature. We synthesized a less reactive dextranol by reducing dextran’s anomeric carbon from an aldehyde to an alcohol. Serum samples dried in a dextranol- based matrix protected the serum proteins from forming high-molecular weight conjugates. The levels of four cancer-related serum biomarkers (prostate specific antigen, neuropilin-1, osteopontin, and metalloproteinase 7) decreased, as measured by immunoassay, when serum samples were stored for one to two weeks in dextran-based matrix. Switching to a dextran-based lyoprotection matrix slightly reduced the damage to osteopontin and completely stopped any detectable damage during storage in the other three biomarkers when for a period of two weeks at 45°C. Dextranol offers a small and easy modification to dextran that significantly improves the molecule’s function as a lyoprotectant by eliminating the potential for damaging protein-polysacharide conjugation.

## Introduction

Room temperature protein preservation is important for food, biologics, purified protein products (e.g. research, cleaning, and industrial enzymes), and clinical samples (1–3). Preservation is required to avoid both chemical (e.g. oxidation) and physical forms of damage (e.g. aggregation) since any damage can reduce therapeutic function, enzymatic activity, flavor, and clinical information value. Many different varieties of excipients have been employed to prevent different forms of damage, one major category being polysaccharides. Large uncharged polymers like polysaccharides protect proteins by molecular crowding, water replacement, and glass formation (4–8).

Dextran has frequently been used as a polysaccharide lyoprotectant in dry protein formulations, mainly due to its high glass transition temperature, which enables room temperature storage (4,9,10). As an inert additive (11), dextran is particularly suitable to be used as a preservative in pharmaceutical products (12–17). As a result, there have been numerous drugs on the market that contain dextran as a preservative (18,19), including biologics (20,21).

Dextran has one drawback, discovered in the food science field in early 1990s; storage of proteins with dextran in low-moisture conditions can facilitate formation of protein-dextran conjugates. (22–25). This reaction has been well documented to occur with various proteins (26–30). Dextran is a branched D-glucose polymer by α-1,6 linkages and α-1,3 linkages at branch points with a single reducing end and multiple non-reducing ends. The conjugation is formed via a Mailard reaction between dextran’s reducing end and protein’s primary amines (N-terminus and/or lysine side chains) leading via a Schiff base to the Amadori product. While these conjugates are mostly shown to form at elevated temperatures (≥50°C) over a time period of days, there are reports of protein-sugar conjugates forming even at lower temperatures, typically over longer time scales. (31–36)

Dextran-protein conjugates are larger and more soluble than un-modified proteins. The size of dextrans, like proteins, covers the low to high kilodalton range and thus, conjugation can easily double or triple the size of the un-modified protein. Like pegylation, dextran conjugation also increases a protein’s solubility (37) and also makes it a better emulsifier (23,38–40). Conjugation of carbohydrates with proteins also causes acidification (lowering of isoelectric point) by removing positive charges from lysine residues (41,42).

Dry storage is an attractive alternative to the typical cryogenic storage for stabilization of proteinaceous biomarkers in clinical biofluid samples like blood serum. Many clinical samples are collected each year and stored in archival biobanks for diagnostic and retrospective biomarker discovery research. The ten largest biobanks collectively house approximately 35 million samples currently (43), stored in mechanical freezers or liquid nitrogen dewars. Cryogenic storage has a number of drawbacks, which could be overcome by room temperature storage of the dried samples; high cost, need to maintain the cold-chain for transportation of samples, and damage to samples due to freeze/thaw are the main ones (44).

Isothermal vitrification of clinical biofluid samples can mitigate the shortcomings of cryogenic storage by allowing storage and handling of samples at room temperature without significant loss of relevant chemical information (2). Isothermal vitrification is achieved by mixing the samples with a lyoprotectant cocktail that increases the glass transition temperature upon water removal to the point that samples form a glassy amorphous solid at or near room temperature. The lyoprotectant matrix that we previously developed (2) uses a mixture of dextran and trehalose with additional low-concentration excipients. The lyoprotectant solution is electrospun into a fiber matrix that, when a biofluid sample is added, simultaneously absorbs the sample as it dissolves and mixes with it. We designed this method to reach a spatially uniform concentration of lyoprotectants in the sample without requiring mixing, which it detrimental to proteins. Overnight drying of the sample-lyoprotectant mix results in a hard, glassy sample that can be stored at room temperature(2).

In our work to preserve clinical serum samples by isothermal vitrification, dextran is essential since it gives the dried samples the high glass transition temperature needed to maintain stability at room temperature. However, conjugation of biomarker proteins with lyoprotectants like dextran could be very detrimental to downstream proteinaceous biomarkers analysis, especially when analytical techniques that are based on specific binding (such as ELISA) are to be utilized to detect activity. In this work, we analyzed the damage to vitrified human serum proteins caused by conjugation with dextran during dried state storage, and we found that replacing dextran in our lyoprotectant formulation with a reduced dextran (dextranol) (**Fig 1**) stabilized serum proteins more effectively during prolonged storage at room temperature as well as at elevated temperatures of 37, and 45°C.

**Figure 1.**
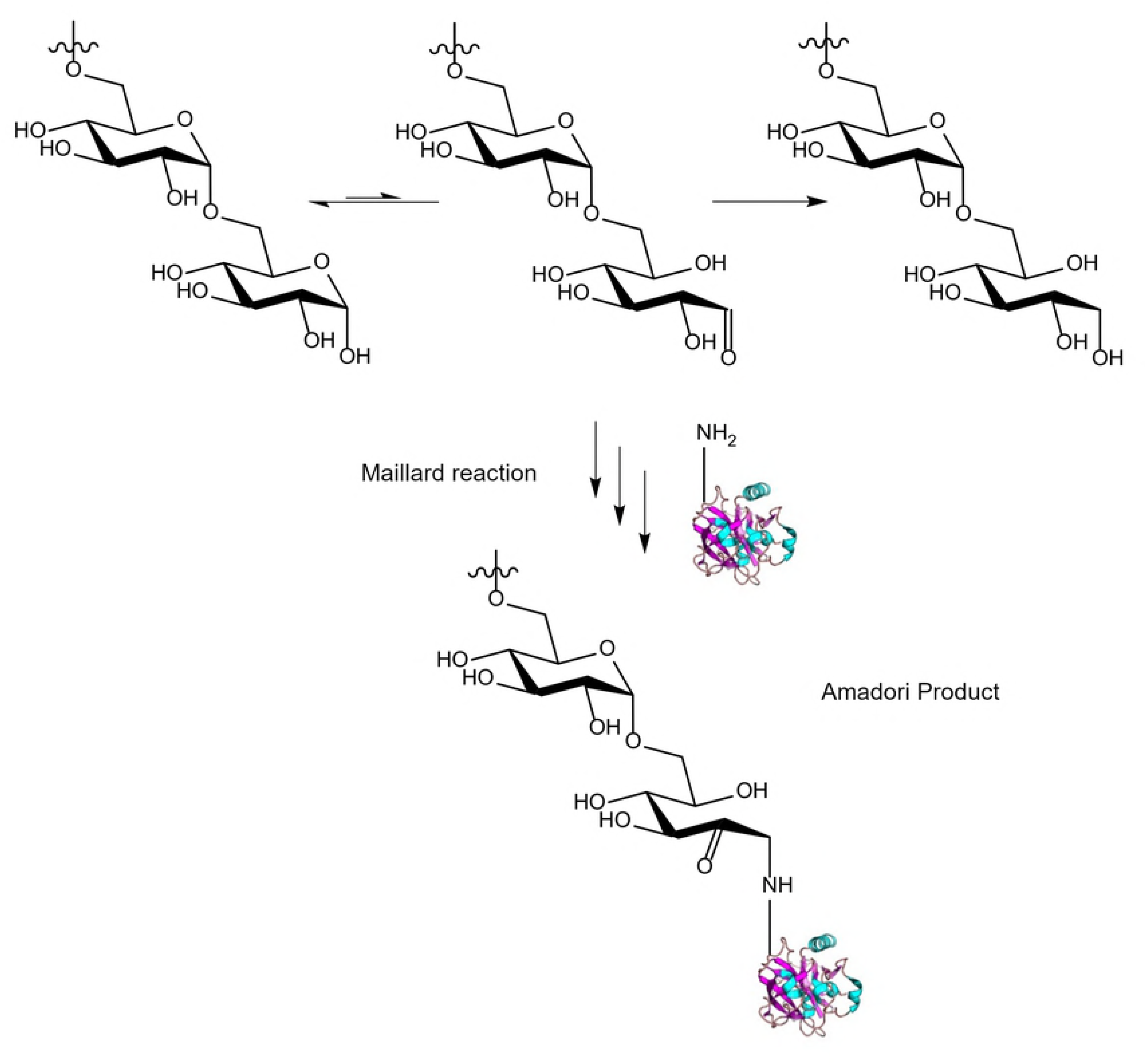
Polysaccharide modification. The cyclic and aldehyde-containing linear forms of the reducing end of a polysaccharide (e.g. dextran) are in equilibrium. The linear form is able to undergo a Maillard reaction with protein amine groups during prolonged storage or upon the addition of heat forming a glycated protein (Schiff base or Amadori Product). Reduction of the polysaccharide aldehyde to an alcohol prevents the Maillard reaction during long term storage of proteins with the polysaccharide.

## Methods

### General

In the experiments conducted here, we used trehalose dihydrate (≥99% purity, Ferro-Pfanstiehl Laboratories, Waukegan, IL), dextran (35-45kDa, Sigma: D1662-500G/lot: SLBT9984), and sodium borohydride (Alfa Aesar: 88983/lot:S09D008). We purchased the other chemicals from Sigma. Anonymized human blood samples were collected from volunteers through the University of Minnesota’s (UMN) Tissue Procurement Facility (TPF) following a UMN Institutional Review Board (IRB) approved protocol (Study Number: 1011E92892). To separate serum, we allowed whole blood to clot for at least 30 minutes and then centrifuged it for 10 min at 2000 RCF. We carefully aspirated the serum (the supernatant) at room temperature and placed it into a new centrifuge tube, taking care not to disturb the cell layer or transfer any cells. We then aliquoted the serum into microcentrifuge tubes for use in experiments.

### Dextranol synthesis

We synthesized dextranol (i.e. reduced dextran) from dextran (from *Leuconostoc mesenteroides* 35-45kDa, Sigma) using a protocol adapted from Paul et al.(45) First, we prepared a solution of 10% w/v dextran in purified water. To this, we added 10 times molar excess of sodium borohydride. We observed mild bubbling as we stirred the solution for 20 hours, after which, we adjusted the pH down to ∼5 using concentrated HCl.

To remove unreacted sodium borohydride and the byproducts, borane and NaCl, we buffer exchanged the solution into water using either dialysis (Fisherbrand 12,000-14,000 MWCO) or spin concentration tubes (Amicon concentrations 3,000 MWCO). Filtration in spin concentrators was very slow, with a flow rate of approximately 5-8 mL/hour at 4000 RCF. We repeated this process until small molecule contaminants were diluted out to less than 1%. Then, we removed water by lyophilization.

### H_1_-NMR

Dextran and dextranol product were each dissolved in DMSO-d_6_ to approximate saturation. We spun tubes to remove the insoluble aggregates and added the supernatant (0.75 mL) to NMR tubes. We collected NMR spectra on Bruker 600 MHz NMR. Disappearance of anomeric proton peaks at 6.7 ppm, and 6.3 ppm, corresponding to the alpha, and beta stereoisomers of the anomeric center, demonstrated the complete reduction to the alcohol (46).

### Production of the nonwoven lyoprotectant matrix by electrospinning

We electrospun fibers to form a dry porous matrix from a lyoprotectant cocktail. The primary components of the cocktail were dextran or dextranol and trehalose. In our previous work, we developed a new dextran-based matrix (V1EX) where five low concentration excipients, 1.5% glycerol (v/v), 1% polyethylene glycol (w/v), 0.1% Tween 20 (v/v), 0.3% gluconic acid (w/v), and 0.2% glucamine (w/v) were incorporated in to the trehalose-dextran cocktail prior to electrospinning. We found the inclusion of these excipients enhanced the stability of the test proteins when desiccated (2). Here we used the same matrix formulation, except where we replaced dextran with dextranol (as described below).

To prepare the lyoprotectant cocktail, we dissolved trehalose (0.4 g/mL) and either dextran or dextranol (1 g/mL) in a solution of low concentration excipients (at concentrations mentioned above). First, we added trehalose to the solution and stirred at 200 RPM for 45 minutes to dissolve it completely. Then, we added dextran or dextranol in three stages, following each with stirring to facilitate dissolution of the solids. We then stirred the mixture overnight (16 hours) at 200 RPM, and at 150 RPM the following day for three hours to eliminate most of the bubbles that formed during mixing. Finally, we allowed the solution to rest for an additional 12 hours at room temperature to ensure total dissolution. We stored the solution at 4°C when not in use.

We electrospun the solution into microfibers across a voltage differential in a controlled environment. We filled 1mL syringes with the lyoprotectant cocktail and affixed a stainless steel 18-gage 0.5“ long blunt-end needle. A multi-channel syringe-pump (NE-1600 multi-syringe pump; New Era Pump Systems, Farmingdale, NY) extruded the solution at a flowrate of 0.03 mL/min. We maintained 50% relative humidity at room temperature in an environment chamber (Electro-tech Systems, Inc., Glenside, PA). We placed the tip of the needle 15 cm away from an aluminum target between which we applied a voltage differential of 15 kV. These conditions were optimized to result in the most uniform electrospun fiber diameter production with optimum inter-fiber distance in the matrix (optimized for capillary adsorption speed vs. dissolution rate) resulting in a well-controlled matrix architecture. After spinning, we dried the matrix in a vacuum chamber overnight to reduce the residual moisture content. We sealed and stored the electrospun matrix in a refrigerator (4°C) until needed.

### Isothermal vitrification and storage

To vitrify serum samples, we aliquoted 50 mg of electrospun fibers into round-bottom screw-top cryogenic vials. To this we added 150 µL of serum. We dried the uncapped tubes in a vacuum chamber (at ≤ -85 kPa pressure) containing Dririte for 24 hours. After this period, we capped the tubes and stored them either at room temperature, or in an incubator set to either 37°C or 45°C for accelerated aging/high-temperature storage experiments. We also prepared biologically matched control samples of serum by freezing and storing the aliquots (without using the lyoprotectant matrix) at −20°C.

To reconstitute the vitrified samples, we added 1.5 mL PBS (Phosphate Buffered Saline) to the dried samples and incubated the tubes for 1 hour with gentle shaking, followed by gentle mixing by pipetting. This resulted in a sample that was 10-fold diluted relative to the original serum. We did not reconstitute the vitrified serum samples to their original volume because this produced a very viscous and difficult to handle solution (due to the presence of lyoprotectant sugars in the solution).

### Differential scanning calorimetry

To determine glass transition temperature of vitrified samples, we collected differential scanning calorimetry (DSC) measurements on a TA Instruments (New Castle, Delaware) DSC Q1000 V9.9 Build 303. A 2-10 mg piece of vitrified serum in either dextran-based or dextranol-matrix was analyzed by ramping the temperature to 150°C at a rate of 10°C/minute after equilibrating at −60°C. Data was collected every second. We analyzed the data using a custom python script that identified the T_g_ (glass transition) as the maximum derivative heat flow of the smoothed data between 20 and 80°C and the T_gonset_(glass transition onset) as the temperature where the heat flow deviated from the linear glass region 10% toward the linear liquid region (each region defined by a linear regression of the most linear 4°C window centered within 5°C lower (glassy region) or higher (liquid region) than the T_g_.

### Gel electrophoresis and staining

We carried out gel electrophoresis using Invitrogen NuPAGE and NativePAGE system (ThermoFisher Scientific, Waltham, MA). For serum analysis, we loaded the equivalent of 0.4 μL of serum (i.e. 4 μL of diluted serum) per well in a 10 well gel. We prepared samples either with NativePAGE buffer for native gel electrophoresis or with LDS sample buffer and the reducing agent for denatured samples, and with LDS sample buffer and the reducing agent for denatured and reduced samples. We boiled both the denatured as well as the denatured and reduced samples for 10 minutes prior to loading on the gel. We ran the gels for 75 minutes at a potential of 150 V.

For general protein stain, we stained native gels using the NativePAGE cathode buffer per manufacturer’s instructions. We stained the denaturing gels for total protein using Imperial Protein Stain (ThermoFisher Scientific). We detected glycoproteins in gels using Pierce Glycoprotein Staining kit (Pierce #24562, lot#SK258276) per manufacturer’s instructions.

## ELISAs

We performed enzyme linked immunoassays (ELISAs) for osteopontin, MMP-7, neuropilin-1, or prostate specific antigen (PSA), following manufacturer’s instructions. We used the following kits: Human Osteopontin ELISA Kit (Abcam, Cambridge, UK, #ab192143), Human Neuropilin-1 ELISA Kit (Abcam #ab227901), Human Total Prostate Specific Antigen ELISA Kit (Abcam #ab188388), and Quantikine ELISA Human Total MMP-7 (R&D Systems Minneapolis, MN, #DMP700). Plates were read in a Tecan Infinite 200 Pro M Nano plate reader at 450nm.

## TCA precipitation

We carried out TCA precipitation by mixing equal parts of resuspended vitrified serum (or the frozen serum control, equivalently diluted after thawing) and 20% TCA (trichloroethanoic acid). After mixing 100 µL of each part, we incubated on ice for 15 minutes. Subsequently, we centrifuged 10 minutes at 10,000 RCF, photographed the tubes, and decanted the supernatant. The pellets were suspended in 400 µL saturated guanidine hydrochloride for protein quantification.

### Protein quantification

Protein was quantified using BCA Protein assay (ThermoFisher Scientific). Standard curve was constructed using bovine serum albumin (BSA) standard. Samples from TCA precipitation were analyzed; both the “soluble” fraction and “precipitate”. Assay was quantified using a Tecan Infinite 200 Pro M Nano plate reader at 562 nm.

## Results

We found that when we used dextran-based lyoprotectant matrix to preserve blood serum by isothermal vitrification, serum began to show signs of damage over time. The damage was detected by the appearance of high-molecular weight smearing seen in gel electrophoresis. While freshly vitrified (and immediately reconstituted) serum appears identical to fresh or frozen serum on gel electrophoresis, after sixteen weeks the smearing becomes very prominent (**Fig 2**), especially when the samples are stored at elevated temperatures. Individual protein bands are less detectable due to decreased intensity and smearing. While smearing is possibly worse in native protein gels (**Fig 2A**), denatured (**Fig 2B**), and denatured & reduced (**Fig 2C**) samples also show smearing. This indicates that the smearing is neither due to non-covalent nor disulfide cross-linked aggregated proteins alone. Non-covalent aggregates would be broken apart by the boiling in LDS (lithium dodecyl sulfate) sample buffer and thus would not be present in the denatured gels. Even if aggregates had disulfide cross-linking, these bonds would be broken by the reducing agent DTT (dithiothreitol) present in the denatured and reduced gel; yet the smearing remains. This indicates that the smearing is due to a large, non-disulfide, covalent modification of proteins.

**Figure 2.**
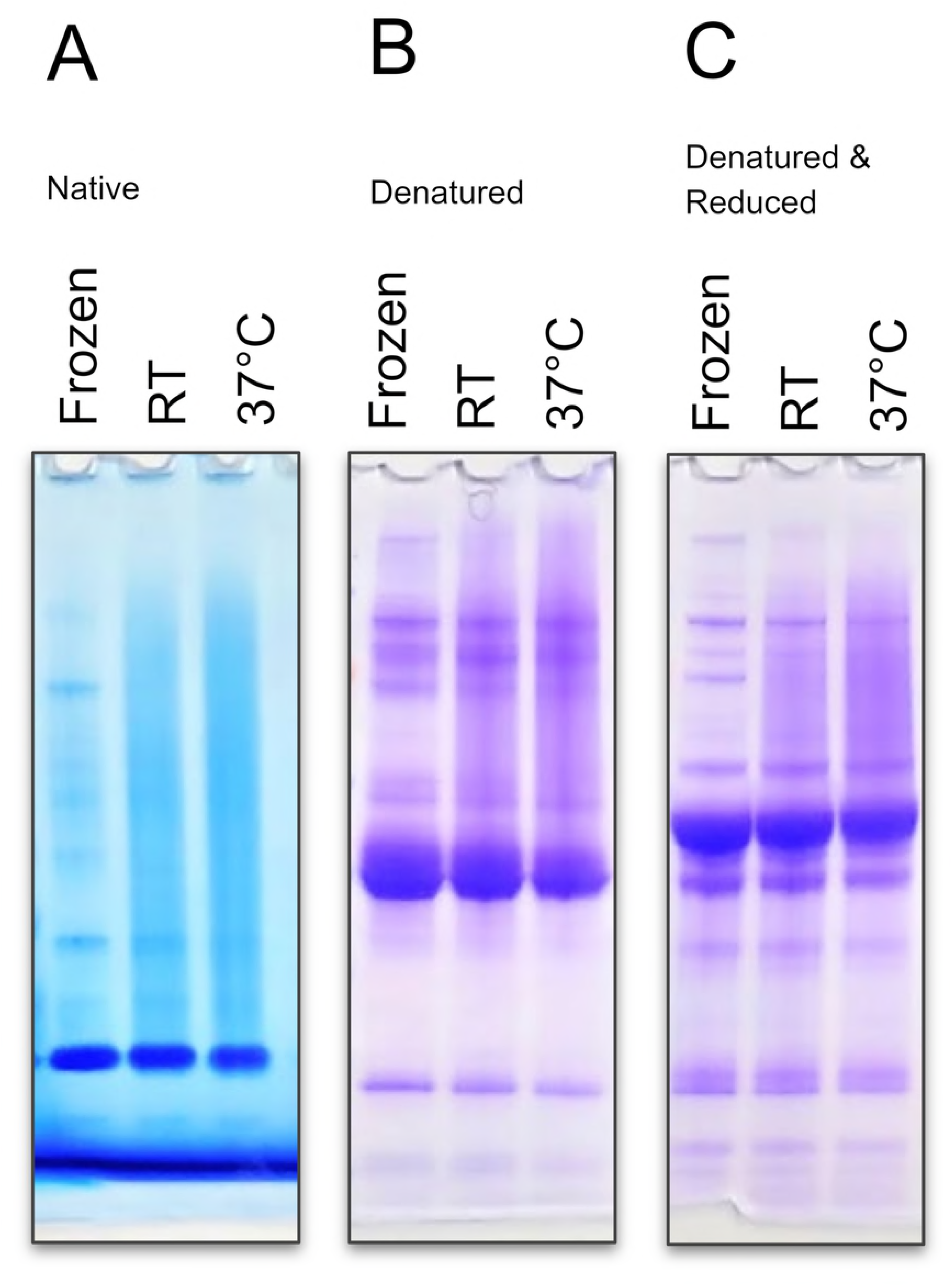
High molecular weight smearing in vitrified samples. Human serum after 16 weeks of storage in the presence of dextran show high molecular weight smearing. Serum was either frozen or vitrified in dextran-based lyoprotectant matrix. Vitrified samples were either stored at room temperature (RT) or 37°C. Vitrified samples were reconstituted in PBS. Gel electrophoresis was carried out using (**A**) native conditions, (**B**) denaturing conditions, or (**C**) denatured & reducing conditions. High molecular weight smears are present in vitrified samples and more pronounced in samples stored at higher temperature. Smearing does not disappear upon denaturation or reduction, showing that it is not due only to denaturation, aggregation, or intramolecular disulfide bond formation, but is due to another covalent modification. This suggests dextran modification of at least some serum proteins.

High molecular weight smearing and glycosylation increased both with storage time and storage temperature and were accompanied by increased solubility. Smearing was faint after only 1 month of storage at room temperature, but increased when either storage temperature was increased to 37°C or storage time was extended to six months (**Fig 3A**). These effects were additive as after six months of storage at elevated temperature the smearing and decrease in individual protein band contrasts was significantly worse than storage either for six months at room temperature or for one month at elevated temperature. Concurrent with the increase in smearing was the increased glycoprotein staining in the high molecular weight region (**Fig 3B**). This indicates that high-molecular weight bodies in the smear are becoming significantly glycosylated. We also noticed that samples that had become more soluble and would not precipitate efficiently using a standard TCA precipitation protocol (**S1 Fig, S1 Table**).

**Figure 3.**
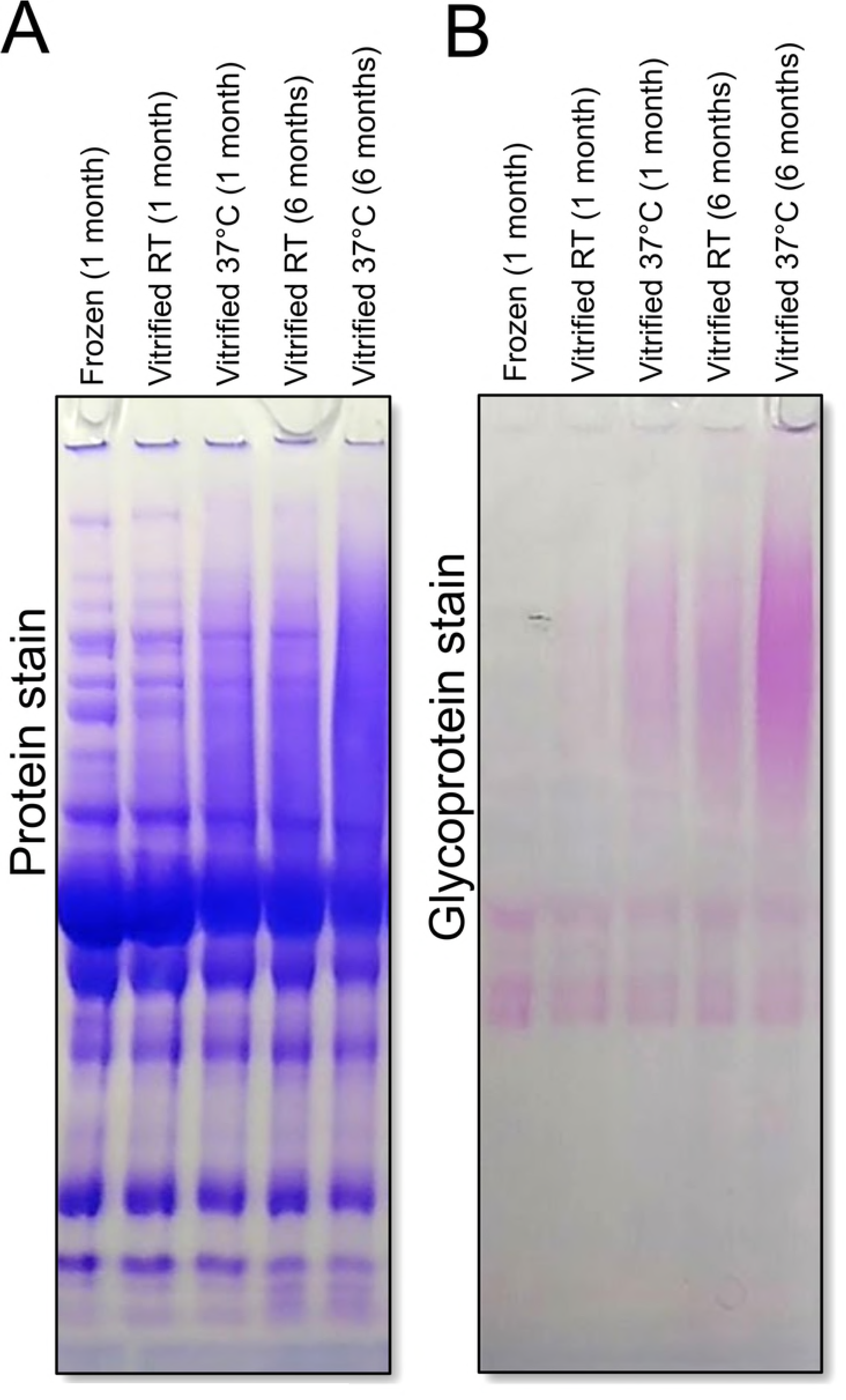
High molecular weight smearing worsens over time and stains positive for glycosylation. Serum samples were vitrified in dextran-based lyoprotectant matrix and stored for 1 or 6 months at either room temperature (RT) or 37°C. Samples were run on SDS-PAGE under denaturing/reducing conditions and stained (**A**) for total protein or (**B**) for glycoprotein. Smearing worsens with higher temperature storage and with increased storage time. Glycoprotein stain indicates high-molecular weight smears are glycosylated; suggesting covalent attachment of dextran to proteins.

Non-reducing dextran, i.e. dextranol, based fibers were able to better preserve serum proteins during extended storage. We reduced the reducing end of dextran from an aldehyde to a more inert alcohol and verified the complete reaction by observing loss of anomeric proton peaks in H_1_-NMR (**S2 Fig**). Dextranol had similar glass-forming properties to dextran (**S3 Fig**). Whereas vitrifying serum in dextran-based lyoprotectant matrix resulted in significant high-molecular weight smearing, serum vitrified in dextranol-based lyoprotectant matrix did not smear (**Figs 4 and S4**). Samples that were frozen, freshly vitrified in dextran-based matrix or dextranol- based matrix were all virtually indistinguishable from fresh, never frozen serum when analyzed by gel electrophoresis under either native (**Fig 4A**) or denaturing conditions (**Fig 4C**). However, after 140 days of storage at 37°C, vitrified samples in dextran-based matrix showed significant smearing, while dextranol-based matrix samples were almost indistinguishable from frozen or fresh serum.

**Figure 4.**
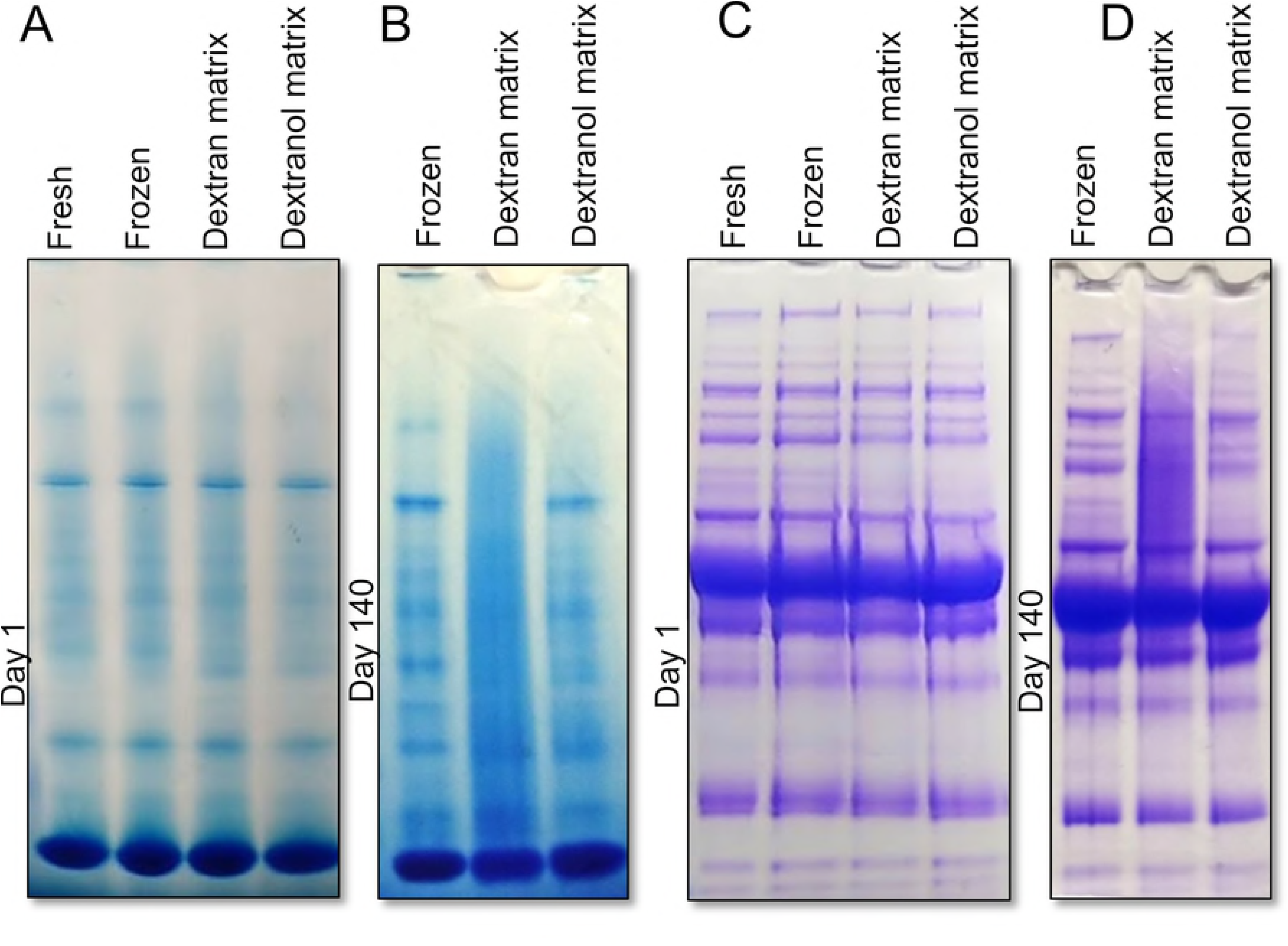
Dextranol-based matrix preserves proteins and prevents smearing compared with dextran-based matrix. Serum samples were either fresh, frozen, or vitrified in either a dextran-based or dextranol-based lyoprotectant matrix. Serum was analyzed (**A** and **C**) immediately after vitrification (Day 1) or (**B** and **D**) after storage for 140 days at 37°C. Vitrified samples were reconstituted in PBS. Gel electrophoresis was carried out under (**A** and **B**) native conditions and (**C** and **D**) denaturing/reducing conditions. After two weeks, smearing is visible in samples preserved in dextran, but not in samples preserved in dextranol. Gels from intermediate days in **S4 Fig.**

Dextranol also protected isothermally vitrified serum and BSA when stored at high temperatures (45°C). The glass transition temperature of the vitrified serum in either dextran or dextranol-based matrix was 50-55°C (**S3 Fig**). Therefore, in all storage experiments, the storage temperatures were kept below 50°C. Storage at higher temperatures allowed us to simulate longer duration storage at room temperature. While decay reactions do not uniformly scale with temperature, we expect a doubling in decay rates every 5-10°C increase (47). Thus 37°C storage corresponded to two to eight times faster aging while 45°C storage corresponded to four to twenty times faster aging. BSA vitrified and stored in dextran-based matrix immediately after drying looked identical to a frozen control, but after storage for 1-2 weeks at 45°C, the band from the main BSA monomer was highly diminished, largely replaced by a high-molecular weight smear (**Fig 5A**). Dextranol-based matrix was able to effectively eliminate this damage, with the sample stored at 45°C for two weeks looking indistinguishable from the freshly vitrified sample and very similar to the frozen aliquot.

**Figure 5.**
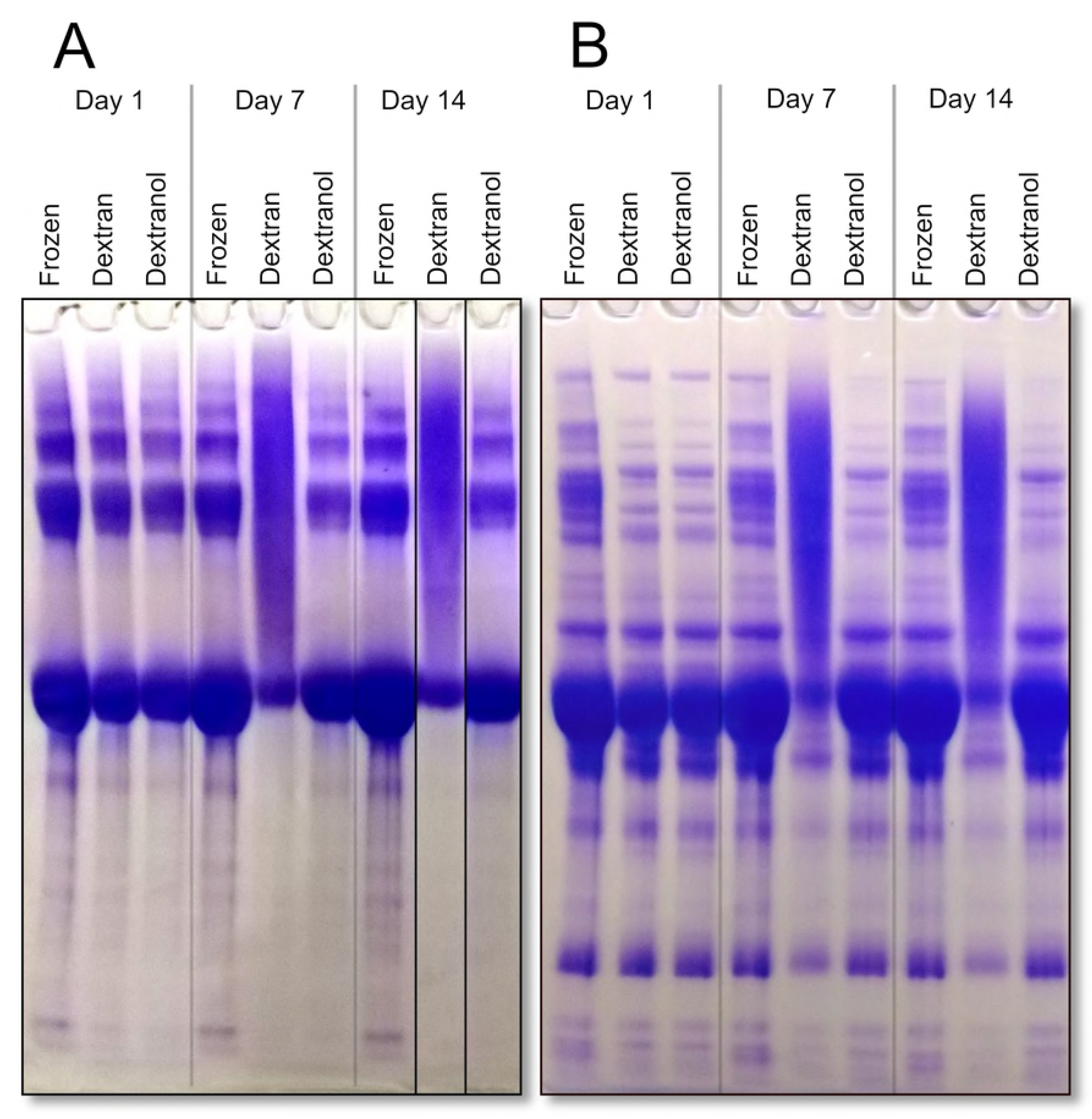
Dextranol protects human serum proteins and BSA when stored at 45° C. Coomassie stained gel showing results of high temperature storage of (**A**) BSA (Bovine serum albumin) and (**B**) human serum. Immediately after vitrification, samples preserved in dextran and dextranol-based lyoprotectant matrices look similar to frozen sample (left). After 7 days (mid), and 14 days (right) BSA stored in dextran-based matrix is not in in the major monomer band but is instead mostly in a high-molecular weight smear. BSA stored in dextranol-based matrix still resembles frozen. Note, it is common to see higher molecular bands in SDS-PAGE of BSA due to irreversible multimer formation.(48,49)

In addition to protecting BSA, the dextranol-based matrix also protected human serum proteins when isothermally vitrified and stored at higher temperature. Most protein bands became largely diminished when the sample was stored for one or two weeks in a dextran-based matrix (**Fig 5B**), and were replaced by a high-molecular weight smear, but storage in dextranol-based matrix protected the proteins. The serum proteins look indistinguishable from freshly vitrified serum after 2 weeks at 45°C, and looked very similar to frozen serum.

In addition to providing overall protection to high abundance serum proteins, and albumin specifically, we also found that dextranol-based lyoprotectant matrix provided much better protection than the dextran-based matrix for four individual proteinaceous biomarkers in the high-temperature stored serum samples. We selected 4 cancer biomarkers to be tested for stability. All four biomarkers that were examined (prostate specific antigen, neuropilin-1, osteopontin, and metalloproteinase 7) showed losses (as measured by ELISA) after one or two weeks of storage at 45°C (**Fig 6** and **S2 Table**). In dextranol-based matrix, PSA (**Fig 6A**) and neuropilin (**Fig 6B**) levels on day 1 were slightly reduced, potentially due to the drying process (by 7 and 8%, respectively), but were stable during seven or fourteen days of storage. MMP-7 levels (**Fig 6D**) in dextranol-based matrices remained slightly above the frozen control throughout this experiment. However, osteopontin levels in the dextranol-based matrices fell in a similar pattern to that seen in dextran-based fibers, although not quite to the same extent.

**Figure 6.**
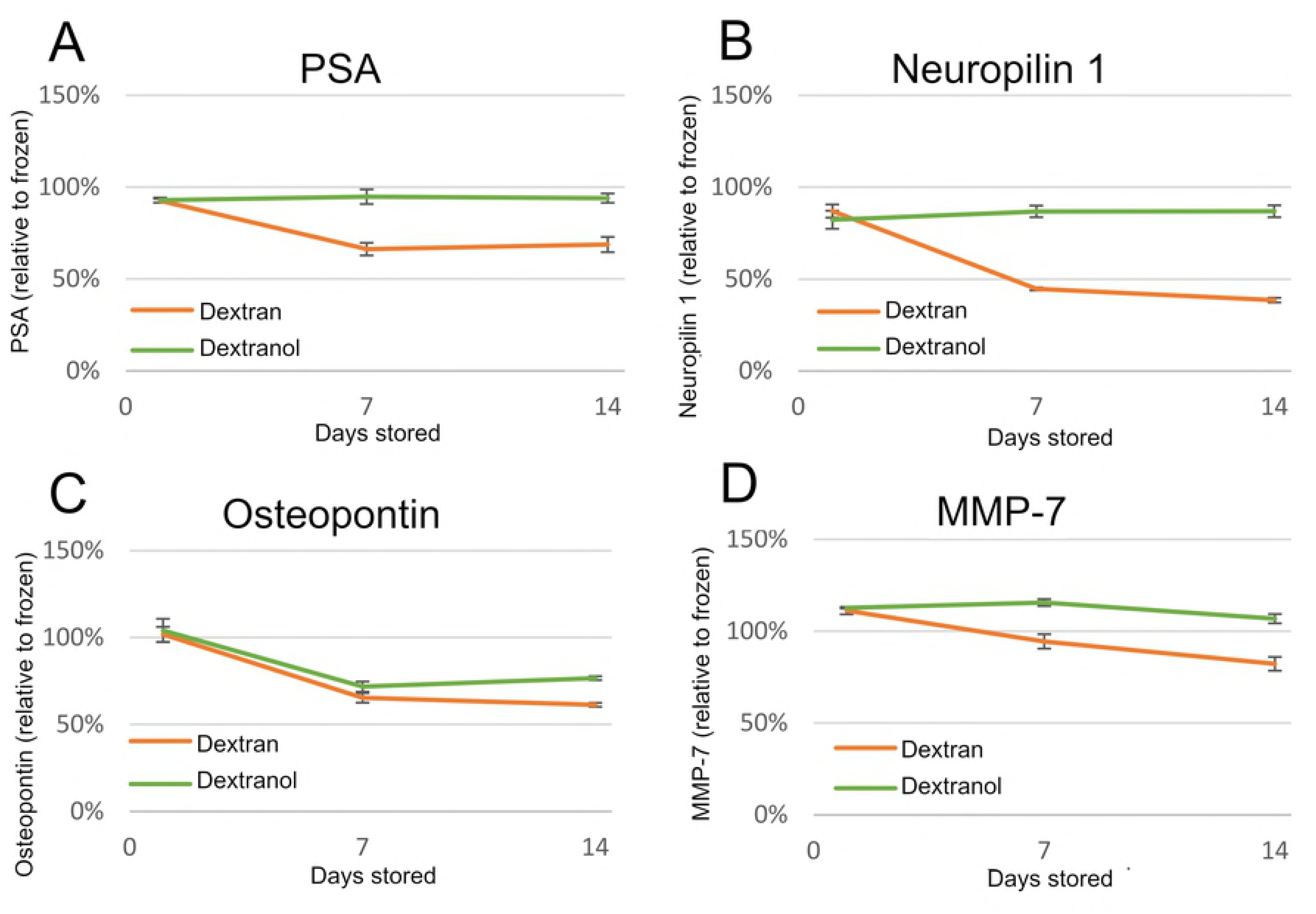
Serum biomarker levels are better retained after storage at 45°C when vitrified in dextranol than dextran. ELISA analysis of biomarker stability in vitrified human serum stored at high temperature (45°C), preserved in either dextran-based (orange) or dextranol-based matrix (green). Four biomarkers examined were (**A**) PSA (prostate specific antigen), (**B**) neuropilin 1, (**C**) osteopontin, and (**D**) MMP-7 (matrix metalloproteinase 7). Serum samples were analyzed immediately after drying (day 1), and one week and two weeks after drying and storage. Values are normalized to biomarker content in frozen control samples. Error bars are standard deviation of three replicates.

## Discussion

Our results show that dextran is not a good lyoprotectant for proteinaceous biomarkers, as it leads to the formation of dextran-protein conjugates, especially in the dried state. Conjugation is likely due to a Maillard reaction forming an Amadori product between the primary amines of a protein (lysine residues and the N termini) and the reducing aldehydes of the dextran chain. While our experiments show this reaction happens to a large extent in a matter of days during storage at higher temperatures (37°C or 45°C), even during room temperature storage, the product is readily apparent after four months. The addition of dextran to proteins was determined to affect their physical properties like solubility as well as their immunogenicity (as seen by decreased reactivity with ELISA antibodies). Decreased immunogenicity results in less accurate immunoassays, which is a problem for detecting biomarkers in stored biospecimens. This damage is concerning since dextran is currently used as a lyoprotectant in a number of pharmaceutical formulations, including formulations for vaccines where decreased immunogenicity would cause reduce efficacy.

Despite the knowledge of the conjugation potential of dextran in food science, researchers in the pharmaceutics filed, in spite of observing protein damage in the presence of dextran, did not identify the mechanism of the damage. Lyophilization research that focused on therapeutic proteins established that formulations containing dextrans cause increase in protein size, increases solubility, and altered acidity, but this was interpreted as aggregation or damage, and not attributed to dextran-protein conjugation. Studies using size exclusion chromatography observe increased protein size after storage with dextran and speculate that this is caused by “soluble aggregation” (50–56). However, size exclusion chromatography cannot distinguish dextran conjugated proteins from dimeric or oligomeric protein “aggregates”. Thus, the “soluble aggregates” reported may instead be dextran conjugates (potentially, in addition to protein self-aggregation). Similarly, studies that used dextran as a lyoprotectant that reported increased solubility (57) and acidity (58) of the preserved proteins could also have observed the effects of dextran conjugation. It’s known that the degree of dextran modification varies between proteins (22,23). This is likely, in part, due to differences in the number and position of surface lysines.

Specifically, Yoshioka et al. characterize dextran mediated damage of lyopholized beta gamma globulin as denaturation and/or aggregation, yet their results are more consistent with dextran-protein conjugation than simple denaturation or aggregation.(50) They conducted experiments with different sizes of dextrans (10kDa-510kDa) and reported that at constant weight fraction in solution, smaller size dextrans (i.e. higher numbers of reactive aldehydes ends per gram of dextran in the solution) caused more damage. They proposed that “the effect of the molecular weight of dextran on the protein stability … could be explained in terms of the parameters obtained by 1H-NMR such as T_mc_ [molecular mobility changing temperature]”. A simpler explanation is that over 50-fold variation in molarity of the reactive aldehyde groups, between the smallest and the largest dextrans, is the main cause. Yoshioka et al. also observed that the increases in sizes of the damaged protein as measured by size-exclusion chromatography correlated with the molecular weight of the dextran; a result that cannot be explained by just denaturation and aggregation, but would be expected if the “damaged proteins” they observed were in fact protein-dextran conjugates.

Qi and Heller, also found evidence for dextran modification of proteins without identifying it as such (12). They report that in liquid state storage, dextran damaged therapeutic peptide insulinotropin. This damage was different from that observed in the absence of dextran and could be prevented by adding certain excipients into the solution (sodium metabisulfate or amino acids) that could react with dextran’s aldehyde group.

Pikal et al. also found evidence for dextran-protein conjugation, which they characterized as soluble “aggregates” (52,53). When they stored human growth hormone freeze-dried in dextran it became larger as measured by size-exclusion chromatography. The increase in the size of the protein in the presence of dextran was greater than when it was lyophilized in its absence or with excipients like glycine, mannitol, hydroxyethyl starch, or trehalose. Interestingly, Pikal et al. suggest that the damage observed when the protein was stored with lactose was due to protein-sugar conjugates formed via a Miallard reaction, yet they didn’t propose the same mechanism for dextran-mediated damage.

Other researchers in pharmaceutical field have looked at intentionally covalently attaching dextran to drug molecules (59). This has been shown to have numerous effects on a drug’s function. Some of the effects are desired; such as improved on-target efficacy (60), reduced toxicity (61), and increased stability (62). However others found detrimental effects from conjugating drugs with dextran, including decreased activity (62) and in one case, a dextran drug conjugate (dextran-doxorubicin) failed in a phase one clinical trial due to higher hepatotoxicity than unmodified doxorubicin (63).

We found that a small chemical modification, converting dextran to dextranol, eliminates this unwanted reaction and better stabilizes human serum proteins during storage. The reduction of dextran’s terminal aldehyde to an alcohol, forming dextranol, ensures inertness. This relatively small chemical modification on a large (∼40 kDa) dextran monomer produced a new lyoprotectant that has all the desirable physical properties of dextran without the potential for the detrimental Maillard reaction. Dextranol has been synthesized before to enhance iron crystal growth and the properties of iron colloids (45,64). However, this communication is the first report on the use of dextranol as an effective lyoprotectant.

Unlike dextran, dextranol-based lyoprotectant matrix we developed was able to protect serum proteins during extended storage with little damage. After 140 days at 37°C or 14 days at 45°C serum proteins stored in dextranol-based matrix showed little sign of degradation by gel electrophoresis, while proteins in dextran-based matrix largely reacted with dextran, visible as high-molecular weight smearing on gels. ELISA analysis of four selected biomarkers showed that detected levels of three of the four biomarkers did not decrease over storage at 45°C when stabilized using the dextranol-based matrix, while levels of all four fell significantly when stored using the dextran-based matrix. Even osteopontin, the one biomarker that did degrade during storage in the dextranol-based matrix, did not lose activity as much as it did in the dextran-based matrix. It is yet unknown what the mechanism of damage was in this case. Osteopontin is frequently cleaved into smaller form, often by thrombin (65). If this were happening in the vitrified sample it may show up as loss of osteopontin since the ELISA antibodies were developed against full-length osteopontin (66). While it is known that inherently disordered proteins like osteopontin (67) are more prone to degradation intracellularly(68), it’s not known if that’s the case extracellularly.

While intentional conjugation of proteins with dextran can have useful applications by improving protein’s solubility and heat stability (27) or as improved food emulsifiers (22), conjugation will inevitably change the physical and chemical properties of the proteins, which can cause loss of function and loss of detection in clinical assays. Other carbohydrates; including lactose, maltodextrin, glucose, and galactomannan; are also known to similarly react with proteins over extended storage times (31–34,69,70). Thus, the use of dextran, or other reducing carbohydrates, as lyoprotectant agents, especially for prolonged times at higher temperatures should be avoided. The modified sugar polymer dextranol is an attractive lyoprotectant since it provides the glass-forming protection of dextran while avoiding the damaging Maillard reaction.

## Acknowledgements

This research was supported by an NIH-NCI grant (R33CA204510).

## Supporting Information

**S1 Figure. TCA precipitation of serum proteins stored 35 days in dextran-based matrix.** Diminished pellet size shows increased solubility somewhat in the sample stored at room temperature (middle tube) and largely in the sample stored at 37°C (right tube) compared to frozen sample without matrix (left tube).

**S1 Table.** **Vitrified serum stored 35 days in dextran-based matrix is not effectively precipitated by TCA.**

|  | Whole<sup>a</sup> | Precipitated<sup>b</sup> |
| --- | --- | --- |
|  | mg/mL<sup>c</sup> | mg/mL<sup>c</sup> |
| Frozen<sup>d</sup> | 59 | 48 |
| Vitrified (RT)<sup>e</sup> | 60 | 46 |
| Vitrified (37° C)<sup>f</sup> | 58 | 12 |

<sup>a</sup> Whole protein concentration after resuspension of vitrified samples and thawing of frozen samples. <sup>b</sup> Protein remaining following resuspension after TCA precipitation. <sup>c</sup> Undiluted serum protein concentration reported.
<sup>d</sup> Serum frozen and stored at -20°C. <sup>e</sup> Serum vitrified in dextran-based matrix and stored at room temperature.
<sup>f</sup> Serum vitrified in dextran-based matrix and stored at 37°C.

**S2 Figure. H_1_-NMR showing complete reaction and disappearance of protons attached to the anomeric carbon.** (**A**) Proton signals at (6.74 and 6.30 ppm) indicate dextran aldehydes, while (**B**) loss of these signals shows that aldehydes were completely reduced to alcohols in dextranol.

**S3 Figure. Tg and Tgon of serum preserved in either dextran and dextranol.** DSC traces of serum samples preserved in either dextran (**A**) or dextranol (**B**) based matrix after 30 days of storage at room temperature. Solid blue line is DSC data, dashed green line is linear glass-transition fit, dashed red line is the linear fit of liquid region (fit to grey-shaded region), dashed cyan line is the liner fit of the glassy region (fit to green-shaded region). The vertical tan line marks the glass transition temperature (T_g_) at 54.7°C (dextran, **A**) and 53.9°C (dextranol, **B**). The vertical gold line marks the glass transition onset temperature (T_gon_) at 53.2°C (dextran, **A**) and 53.1°C (dextranol, **B**).

**S4 Figure. Dextranol preserves proteins and prevents smearing compared with dextran.** Serum samples were either fresh, frozen, or vitrified in either a dextran-based or dextranol-based lyoprotectant matrix. Serum was analyzed after 1, 7, 14, 28, 60, and 140 days at 37°C (data from days 1 and 140 in **Fig 4**). Vitrified samples were reconstituted in PBS. Gel electrophoresis was carried out under both native conditions (**A**) and denaturing/reducing (**B**) conditions. A duplicate denatured/reduced gel from the sixty day old sample was also stained for glycoproteins (**C**). After two weeks, smearing is visible in samples preserved in dextran, but not in samples preserved in dextranol.

**S2 Table.**
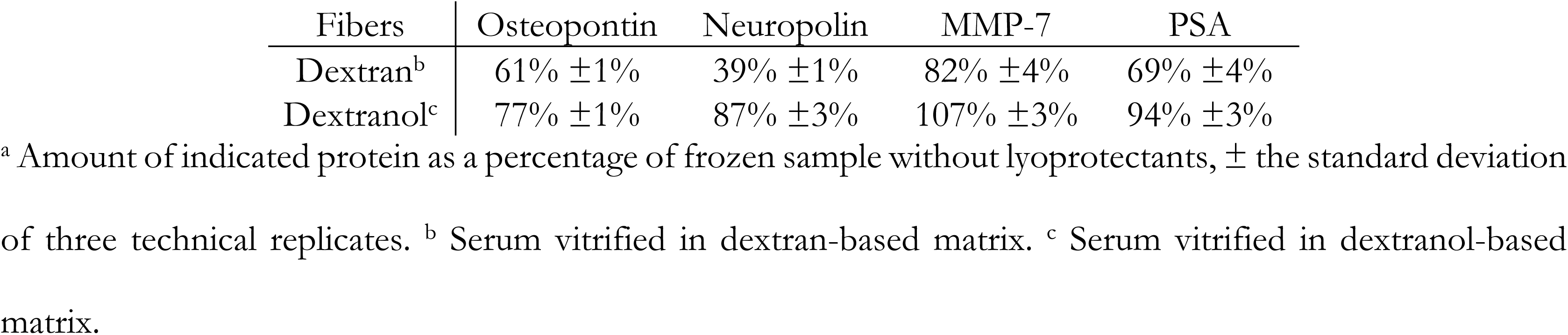
**Biomarker preservation after 14 days storage of vitrified samples at 45°Ca**

| Fibers | Osteopontin | Neuropilin | MMP-7 | PSA |
| --- | --- | --- | --- | --- |
| Dextran<sup>b</sup> | 61% ±1% | 39% ±1% | 82% ±4% | 69% ±4% |
| Dextranol<sup>c</sup> | 77% ±1% | 87% ±3% | 107% ±3% | 94% ±3% |

<sup>a</sup> Amount of indicated protein as a percentage of frozen sample without lyoprotectants, ± the standard deviation of three technical replicates. <sup>b</sup> Serum vitrified in dextran-based matrix. <sup>c</sup> Serum vitrified in dextranol-based matrix.

## References

1. Wang W. Advanced protein formulations. Protein Sci Publ Protein Soc. 2015 Jul;24(7):1031–9.

2. Solivio MJ, Less R, Rynes ML, Kramer M, Aksan A. Adsorbing/dissolving Lyoprotectant Matrix Technology for Non-cryogenic Storage of Archival Human Sera. Sci Rep. 2016;6:24186.

3. Boye JI, Barbana C. 3 Protein Processing in Food and Bioproduct Manufacturing and Techniques for Analysis. Food Ind Bioprod Bioprocess. 2012;85.

4. Aksan A, Toner M. Isothermal Desiccation and Vitrification Kinetics of Trehalose- Dextran Solutions. Langmuir. 2004;20(13):5521–5529.

5. Minton AP. Quantitative Assessment of the Relative Contributions of Steric Repulsion and Chemical Interactions to Macromolecular Crowding. Biopolymers. 2013 Apr;99(4):239–44.

6. Sasahara K, McPhie P, Minton AP. Effect of dextran on protein stability and conformation attributed to macromolecular crowding. J Mol Biol. 2003;326(4):1227–1237.

7. Aksan A, Irimia D, He X, Toner M. Desiccation kinetics of biopreservation solutions in microchannels. J Appl Phys. 2006;99(6):064703.

8. Ragoonanan V, Aksan A. Heterogeneity in Desiccated Solutions: Implications for Biostabilization. Biophys J. 2008 Mar 15;94(6):2212–27.

9. Yoshioka S, Miyazaki T, Aso Y. β-Relaxation of Insulin Molecule in Lyophilized Formulations Containing Trehalose or Dextran as a Determinant of Chemical Reactivity. Pharm Res. 2006 May 1;23(5):961–6.

10. Allison SD, Manning MC, Randolph TW, Middleton K, Davis A, Carpenter JF. Optimization of Storage Stability of Lyophilized Actin Using Combinations of Disaccharides and Dextran. J Pharm Sci. 2000 Feb 1;89(2):199–214.

11. FDA Center for Drug Evaluation and Research. Inactive Ingredient Search for Approved Drug Products [Internet]. 2018 [cited 2018 Jun 28]. Available from: https://www.accessdata.fda.gov/scripts/cder/iig/index.cfm

12. Qi H, Heller DL. Stability and Stabilization of Insulinotropin in a Dextran Formulation. PDA J Pharm Sci Technol. 1995 Nov 1;49(6):289–93.

13. Yuan W, Geng Y, Wu F, Liu Y, Guo M, Zhao H, et al. Preparation of polysaccharide glassy microparticles with stabilization of proteins. Int J Pharm. 2009;366(1–2):154–159.

14. Nakade S, Komaba J, Ohno T, Kitagawa J, Furukawa K, Miyata Y. Bioequivalence study of two limaprost alfadex 5 microg tablets in healthy subjects: moisture-resistant tablet (dextran formulation) versus standard tablet (lactose formulation). Int J Clin Pharmacol Ther. 2008 Jan;46(1):42–7.

15. Parkash V, Maan S, Deepika, Yadav SK, Hemlata, Jogpal V. Fast disintegrating tablets: Opportunity in drug delivery system. J Adv Pharm Technol Res. 2011;2(4):223–35.

16. Wilkhu JS, McNeil SE, Anderson DE, Kirchmeier M, Perrie Y. Development of a solid dosage platform for the oral delivery of bilayer vesicles. Eur J Pharm Sci. 2017 Oct 15;108:71–7.

17. Sekiya N, Abe N, Ishikawa H, Inoue Y, Yamamoto M, Takeda K. Improved stability of OPALMON tablets under humid conditions II: stability in one-dose package. Iryo Yakugaku. 2006;32:482–488.

18. Bristol-Myers Squibb. Etopophos: Highlights of Prescribing Information [Internet]. Bristol-Myers Squibb; 2017. Available from: packageinserts.bms.com/pi/pi_etopophos.pdf

19. Mercury Pharmaceuticals. Patient Information Leaflet: Liothyronine Sodium 20 micrograms Powder for Solution for Injection [Internet]. Mercury Pharmaceuticals; 2013. Available from: http://www.medicines.org.uk/emc/files/pil.2805.pdf

20. Wyeth Pharmaceuticals Inc. Mylotarg: Highlights of Prescribing Information [Internet]. Wyeth Pharmaceuticals Inc.; 2018. Available from: https://www.mylotarg.com/

21. GlaxoSmithKline. Rotarix: Highlights of Prescribing Information [Internet]. GlaxoSmithKline; 2016. Available from: http://www.gsksource.com/pharma/content/dam/GlaxoSmithKline/US/en/Prescribing_Information/Rotarix/pdf/ROTARIX-PI-PIL.PDF

22. Kato A, Sasaki Y, Furuta R, Kobayashi K. Functional protein-polysaccharide conjugate prepared by controlled dry-heating of ovalbumin-dextran mixtures. Agric Biol Chem. 1990;54(1):107–112.

23. Kato A, Mifuru R, Matsudomi N, Kobayashi K. Functional casein-poly saccharide conjugates prepared by controlled dry heating. Biosci Biotechnol Biochem. 1992;56(4):567–571.

24. Nakamura S, Kato A, Kobayashi K. New antimicrobial characteristics of lysozyme-dextran conjugate. J Agric Food Chem. 1991;39(4):647–650.

25. Nakamura S, Kato A, Kobayashi K. Novel bifunctional lysozyme–dextran conjugate that acts on both Gram-negative and Gram-positive bacteria. Agric Biol Chem. 1990;54(11):3057–3059.

26. de Oliveira FC, Coimbra JS dos R, de Oliveira EB, Zuñiga ADG, Rojas EEG. Food protein- polysaccharide conjugates obtained via the maillard reaction: A review. Crit Rev Food Sci Nutr. 2016;56(7):1108–1125.

27. Jiménez-Castaño L, Villamiel M, López-Fandiño R. Glycosylation of individual whey proteins by Maillard reaction using dextran of different molecular mass. Food Hydrocoll. 2007;21(3):433–443.

28. Jiménez-Castaño L, Villamiel M, Martín-Álvarez PJ, Olano A, López-Fandiño R. Effect of the dry-heating conditions on the glycosylation of β-lactoglobulinv with dextran through the Maillard reaction. Food Hydrocoll. 2005;19(5):831–837.

29. Spotti MJ, Perduca MJ, Piagentini A, Santiago LG, Rubiolo AC, Carrara CR. Does dextran molecular weight affect the mechanical properties of whey protein/dextran conjugate gels? Food Hydrocoll. 2013;32(1):204–210.

30. Liu Y, Zhao G, Zhao M, Ren J, Yang B. Improvement of functional properties of peanut protein isolate by conjugation with dextran through Maillard reaction. Food Chem. 2012;131(3):901–906.

31. Bell LN, Touma DE, White KL, Chen Y-H. Glycine loss and Maillard browning as related to the glass transition in a model food system. J Food Sci. 1998;63(4):625–628.

32. Intipunya P, Bhandari BR. Chemical deterioration and physical instability of food powders. In: Chemical Deterioration and Physical Instability of Food and Beverages. Elsevier; 2010. p. 663–700.

33. Baptista JA, Carvalho RC. Indirect determination of Amadori compounds in milk-based products by HPLC/ELSD/UV as an index of protein deterioration. Food Res Int. 2004;37(8):739–747.

34. Kim MN, Saltmarch M, Labuza TP. Non-Enzymatic Browning of Hygroscopic Whey Powders in Open Versu Sealed Pouches. J Food Process Preserv. 1981;5(1):49–57.

35. Hrynets Y, Ndagijimana M, Betti M. Non-enzymatic glycation of natural actomyosin (NAM) with glucosamine in a liquid system at moderate temperatures. Food Chem. 2013;139(1–4):1062–1072.

36. Gaucher I, Mollé D, Gagnaire V, Gaucheron F. Effects of storage temperature on physico-chemical characteristics of semi-skimmed UHT milk. Food Hydrocoll. 2008;22(1):130–143.

37. Jiménez-Castaño L, López-Fandiño R, Olano A, Villamiel M. Study on β-lactoglobulin glycosylation with dextran: effect on solubility and heat stability. Food Chem. 2005;93(4):689–695.

38. Dickinson E, Galazka VB. Emulsion stabilization by ionic and covalent complexes of β-lactoglobulin with polysaccharides. Food Hydrocoll. 1991;5(3):281–296.

39. Admassu H, Zhao W, Yang R, Gasmalla MA, Alsir El. Stabilizing Food Emulsions By Protein–Polysaccharide Conjugates Of Maillard Reaction-A. 2015;3(8):103–8.

40. Ho Y-T, Ishizaki S, Tanaka M. Improving emulsifying activity of ε-polylysine by conjugation with dextran through the Maillard reaction. Food Chem. 2000;68(4):449–455.

41. Luthra M, Balasubramanian D. Nonenzymatic glycation alters protein structure and stability. A study of two eye lens crystallins. J Biol Chem. 1993;268(24):18119–18127.

42. Nacka F, Chobert J-M, Burova T, Léonil J, Haertlé T. Induction of new physicochemical and functional properties by the glycosylation of whey proteins. J Protein Chem. 1998;17(5):495–503.

43. Orchard-Webb D. 10 Largest Biobanks in the World [Internet]. Biobanking.com. 2018 [cited 2018 Sep 28]. Available from: https://www.biobanking.com/10-largest-biobanks-in-the-world/

44. Bortolin RC, Gasparotto J, Vargas AR, da Silva Morrone M, Kunzler A, Henkin BS, et al. Effects of Freeze-Thaw and Storage on Enzymatic Activities, Protein Oxidative Damage, and Immunocontent of the Blood, Liver, and Brain of Rats. Biopreservation Biobanking. 2017 Jun;15(3):182–90.

45. Paul KG, Frigo TB, Groman JY, Groman EV. Synthesis of ultrasmall superparamagnetic iron oxides using reduced polysaccharides. Bioconjug Chem. 2004;15(2):394–401.

46. Kuang H, Wu Y, Zhang Z, Li J, Chen X, Xie Z, et al. Double pH-responsive supramolecular copolymer micelles based on the complementary multiple hydrogen bonds of nucleobases and acetalated dextran for drug delivery. Polym Chem. 2015;6(19):3625–3633.

47. Michalski S. Double the life for each five-degree drop, more than double the life for each halving of relative humidity. In: Preprints of 13th Meeting of ICOM-CC. p. 66–72.

48. Brahma A, Mandal C, Bhattacharyya D. Characterization of a dimeric unfolding intermediate of bovine serum albumin under mildly acidic condition. Biochim Biophys Acta BBA - Proteins Proteomics. 2005 Aug 10;1751(2):159–69.

49. Wong C-Y, Chung L-H, Lin S, Chan DS-H, Leung C-H, Ma D-L. A Ruthenium(II) Complex Supported by Trithiacyclononane and Aromatic Diimine Ligand as Luminescent Switch-On Probe for Biomolecule Detection and Protein Staining. Sci Rep. 2014 Nov 20;4:7136.

50. Yoshioka S, Aso Y, Kojima S. Dependence of the Molecular Mobility and Protein Stability of Freeze- Dried γ-Globulin Formulations on the Molecular Weight of Dextran. Pharm Res. 1997 Jun 1;14(6):736–41.

51. Kreilgaard L, Frokjaer S, Flink JM, Randolph TW, Carpenter JF. Effects of Additives on the Stability of Recombinant Human Factor XIII during Freeze-Drying and Storage in the Dried Solid. Arch Biochem Biophys. 1998 Dec 1;360(1):121–34.

52. Pikal MJ, Dellerman KM, Roy ML, Riggin RM. The effects of formulation variables on the stability of freeze-dried human growth hormone. Pharm Res. 1991;8(4):427–436.

53. Pikal MJ. 8 - Mechanisms of Protein Stabilization During Freeze-Drying Storage: The Relative Importance of Thermodynamic Stabilization and Glassy State Relaxation Dynamics. In: Rey L, May JC, editors. Freeze-Drying/Lyophilization of Pharmaceutical and Biological Products, Third Edition. 3rd ed. CRC Press; 2004. p. 198–232.

54. Schüle S, Schulz-Fademrecht T, Garidel P, Bechtold-Peters K, Frie\s s W. Stabilization of IgG1 in spray- dried powders for inhalation. Eur J Pharm Biopharm. 2008;69(3):793–807.

55. Lueckel B, Helk B, Bodmer D, Leuenberger H. Effects of Formulation and Process Variables on the Aggregation of Freeze-Dried Interleukin-6 (IL-6) After Lyophilization and on Storage. Pharm Dev Technol. 1998 Jan 1;3(3):337–46.

56. Kreilgaard L, Frokjaer S, Flink JM, Randolph TW, Carpenter JF. Effects of additives on the stability of Humicola lanuginosa lipase during freeze-drying and storage in the dried solid. J Pharm Sci. 1999;88(3):281–90.

57. Costantino HR, Griebenow K, Mishra P, Langer R, Klibanov AM. Fourier-transform infrared spectroscopic investigation of protein stability in the lyophilized form. Biochim Biophys Acta BBA - Protein Struct Mol Enzymol. 1995 Nov 15;1253(1):69–74.

58. Ressing ME, Jiskoot W, Talsma H, Van Ingen CW, Beuvery EC, Crommelin DJ. The influence of sucrose, dextran, and hydroxypropyl-β-cyclodextrin as lyoprotectants for a freeze-dried mouse IgG2a monoclonal antibody (MN12). Pharm Res. 1992;9(2):266–270.

59. Takakura Y, Takagi A, Hashida M, Sezaki H. Disposition and tumor localization of mitomycin C–dextran conjugates in mice. Pharm Res. 1987;4(4):293–300.

60. Mitra S, Gaur U, Ghosh PC, Maitra AN. Tumour targeted delivery of encapsulated dextran–doxorubicin conjugate using chitosan nanoparticles as carrier. J Controlled Release. 2001 Jul 6;74(1):317–23.

61. Levi-Schaffer F, Bernstein A, Meshorer A, Arnon R. Reduced toxicity of daunorubicin by conjugation to dextran. Cancer Treat Rep. 1982 Jan;66(1):107–14.

62. Takakura Y, Kaneko Y, Fujita T, Hashida M, Maeda H, Sezaki H. Control of pharmaceutical properties of soybean trypsin inhibitor by conjugation with dextran I: Synthesis and characterization. J Pharm Sci. 1989;78(2):117–121.

63. Danhauser-Riedl S, Hausmann E, Schick H-D, Bender R, Dietzfelbinger H, Rastetter J, et al. Phase I clinical and pharmacokinetic trial of dextran conjugated doxorubicin (AD-70, DOX-OXD). Invest New Drugs. 1993;11(2–3):187–95.

64. Floramo NA. Iron hydrogenated dextran [Internet]. US3022221A, 1962 [cited 2018 Sep 27]. Available from: https://patents.google.com/patent/US3022221A/en?oq=DK+117%2c730

65. Lanteri P, Lombardi G, Colombini A, Grasso D, Banfi G. Stability of osteopontin in plasma and serum. Clin Chem Lab Med [Internet]. 2012 Jan 1 [cited 2018 Nov 6];50(11). Available from: https://www.degruyter.com/view/j/cclm.2012.50.issue-11/cclm-2012-0177/cclm-2012-0177.xml

66. Human Osteopontin ELISA Kit (ab100618) | Abcam [Internet]. [cited 2018 Nov 6]. Available from: https://www.abcam.com/human-osteopontin-elisa-kit-ab100618.html?productWallTab=ShowAll

67. Fisher LW, Torchia DA, Fohr B, Young MF, Fedarko NS. Flexible Structures of SIBLING Proteins, Bone Sialoprotein, and Osteopontin. Biochem Biophys Res Commun. 2001 Jan 19;280(2):460–5.

68. Tompa P, Prilusky J, Silman I, Sussman JL. Structural disorder serves as a weak signal for intracellular protein degradation. Proteins Struct Funct Bioinforma. 2008 May 1;71(2):903–9.

69. Evangelisti F, Calcagno C, Zunin P. Relationship Between Blocked Lysine and Carbohydrate Composition of Infant Formulas. J Food Sci. 1994 Mar;59(2):335–7.

70. Babiker E fadil E, Hiroyuki A, Matsudomi N, Iwata H, Ogawa T, Bando N, et al. Effect of polysaccharide conjugation or transglutaminase treatment on the allergenicity and functional properties of soy protein. J Agric Food Chem. 1998;46(3):866–871.

